# FCS-Like Zinc finger 14 (FLZ14) mediates the crosstalk between TORC and SnRK1 in response to sugar availability

**DOI:** 10.1101/2023.08.14.553285

**Authors:** Anthony Artins, Reynel Urrea-Castellanos, Marcos Martin-Sánchez, Thiago Alexandre Moraes, Alisdair R. Fernie, Akiko Satake, Camila Caldana

## Abstract

Matching resource availability to growth is crucial for plant fitness. We identified that FCS- Like Zinc finger 14 (FLZ14) gauges carbon (C) status to adjust growth by inhibiting the TARGET OF RAPAMYCIN (TOR) activity in short days via REGULATORY- ASSOCIATED PROTEIN OF TARGET OF RAPAMYCIN 1B (RAPTOR1B). Besides being transcriptionally induced by sugars, the amplitude of *FLZ14* induction correlates to the diel sugar status, indicating that *FLZ14* responses to sugar oscillations *in planta.* Genetic evidence elucidated that FLZ14 is involved in the crosstalk between TORC and SUCROSE NON-FERMENTING RELATED KINASE 1 (SnRK1). In short photoperiods, *FLZ14* and *KIN10* genetically interact to set the pace of TORC activity. By contrast, under long day photoperiods, *FLZ14* and *RAPTOR1B* genetically interact, most likely to downregulate SnRK1 signalling. This report highlights the function of a sugar-responsive gene in mediating the crosstalk between TORC and SnRK1 to fine-tune growth.

## Introduction

The sessile nature of plants, their source and sink tissues, and their multicellularity require strict coordination of physiological responses to the resource availability to sustain growth. This process is orchestrated by a complex regulatory network in which the TARGET OF RAPAMYCIN (TOR) kinase is one of the central hubs. In plants, TOR assembles a single protein complex (TORC) with LETHAL WITH SEC THIRTEEN PROTEIN 8 (LST8) and REGULATORY-ASSOCIATED PROTEIN OF TOR (RAPTOR) activated by high energy and nutrient status. RAPTOR acts as a scaffolding protein to recruit substrates such as ribosomal S6 Kinase (S6K) and 4E-binding protein1 (4E-BP1) for phosphorylation by the TOR kinase (Nojima et al., 2003; Schalm et al., 2003). In Arabidopsis, this regulatory protein is encoded by two genes, *AtRAPTOR1A* and *AtRAPTOR1B*; however, only mutations in the latter have a significant impact on plant growth and metabolism (Anderson et al., 2005; Salem et al., 2018). Evidence suggests that RAPTOR1B conveys signals to TOR, intertwining with other branches of the growth-mediated regulatory network, such as SUCROSE-NON-FERMENTING 1 (SNF1)-RELATED KINASE 1 (SnRK1). Under energy and sugar limitation, SnRK1 interacts with and phosphorylates RAPTOR1B, inactivating TORC (Nukarinen et al., 2016).

The starvation-induced genes *FCS-LIKE ZINC FINGER (FLZ) 6* and *10* were shown to prevent the hyper-activation of SnRK1, thereby alleviating TORC repression (Jamsheer et al., 2018). Under unexpected C starvation ATG8, a component of the autophagy conjugation pathway, degrades AIM-containing FLZ proteins, such as FLZ14, derepressing SnRK1 activity (Yang et al., 2023). The plant-specific FLZ family has 18 members, each responding to a certain environmental cue such as nutrient availability and abiotic stress or endogenous hormone cues (Nietzsche et al., 2014), placing these proteins as interesting candidates for providing signal specificity to the TORC-SnRK1 signalling network. While all FLZ members interact with SnRK1 (Jamsheer and Laxmi, 2014; Nietzsche et al., 2014), only FLZ9 and FLZ8 were confirmed to interact with RAPTOR1B so far (Nietzsche et al., 2016). Under favourable growth conditions, FLZ8 bridges SnRK1α1 and RAPTOR1B, thereby downregulating TOR signalling (Jamsheer et al., 2022). These studies indicate that SnRK1 and TOR are not only antagonistic pathways but are fine-tuned to control metabolic homeostasis to adjust growth according to resource availability.

Here, we identified the interaction between RAPTOR1B and FLZ14, which was previously shown to respond to sugar levels (Jamsheer K and Laxmi, 2015). In contrast to previous work, we focused on the consequences of this interaction in the regulation of growth during the diel cycle. The alternation of light and dark periods imposes fluctuations in sugar levels, with the photoperiod length dictating carbon (C) partitioning, allocation, and use for growth and development (Gibon et al., 2009; Sulpice et al., 2009, 2014). The longer the photoperiod, the greater the C availability allocated for growth (Yazdanbakhsh et al., 2011; Sulpice et al., 2014; Apelt et al., 2017). By contrast, low C availability imposed by short days (SD) leads to larger oscillations in the sugar levels between day and night (Gibon et al., 2004, 2009; Mengin et al., 2017). In this scenario, we found that FLZ14 reduces the pace of TORC activity, acting as a growth repressor. This negative regulation appears to occur through the action of KIN10, the predominant kinase subunit of the SnRK1 complex. Conversely, the dampened fluctuations in sugar levels in long days (LD) favour the genetic interaction between *FLZ14* and *RAPTOR1B* to promote growth. This regulation between TORC and SnRK1 mediated by FLZ14 allows the maintenance of metabolic homeostasis, optimising plant growth by matching the availability of resources.

## Results

### FLZ14 interacts with RAPTOR1B, affecting growth in a photoperiod-dependent manner

We performed a Yeast-2 Hybrid (Y2H) screening using the regulatory subunit of TORC, *RAPTOR1B*, as a bait to identify new players involved in the sugar signaling network. We found the sugar-responsive gene *FCS-Like Zinc Finger (FLZ)14* (AT5G20700) (Nietzsche et al., 2014; Jamsheer K and Laxmi, 2015) as a potential interacting partner of RAPTOR1B, together with 85 other candidates **(Supplemental Table 1)**, including known interactors such as the RIBOSOMAL S6 KINASE 1 (S6K1) (Hara et al., 2002; Kim et al., 2002; Nojima et al., 2003; Mahfouz et al., 2006) and MEDIATOR OF ABA-REGULATED DORMANCY 1 (MARD1/FLZ9) (Nietzsche et al., 2016). The physical interaction between RAPTOR1B and FLZ14 was confirmed by an independent targeted Y2H assay **(Supplemental Figure 1A)**. To validate this interaction *in planta*, a co-immunoprecipitation (Co-IP) was performed using RAPTOR1B-6xMyc and FLZ14-3xHA tagged proteins, and antibodies that could recognise either Myc or HA. FLZ14-3xHA was co-immunoprecipitated with RAPTOR1B-6xMyc. Likewise, RAPTOR1B-6xMyc was detected by immunoprecipitating FLZ14-3xHA, while no signal was obtained when only a single construct was infiltrated into tobacco leaves **(Figure 1A)**. Similar results were found using bimolecular fluorescence complementation (BiFC) assays **(Supplemental Figure 1B)**. Co-infiltration of either *nYFP:RAPTOR1B_cDNA_ / cYFP:FLZ14_cDNA_* or *nYFP:RAPTOR1B_cDNA_ / FLZ14_cDNA_:cYFP* constructs into tobacco leaves unravelled that the interaction takes place in both epidermal and stomatal guard cells **(Supplemental Figure 1B)**. By contrast, no interaction was observed between FLZ6 and RAPTOR1B (*nYFP:RAPTOR1B_cDNA_ / FLZ6_cDNA_:cYFP*), used as a negative control **(Supplemental Figure 1B)**. Taken together, these results confirm FLZ14 as a new interactor of RAPTOR1B.

**Figure 1.**
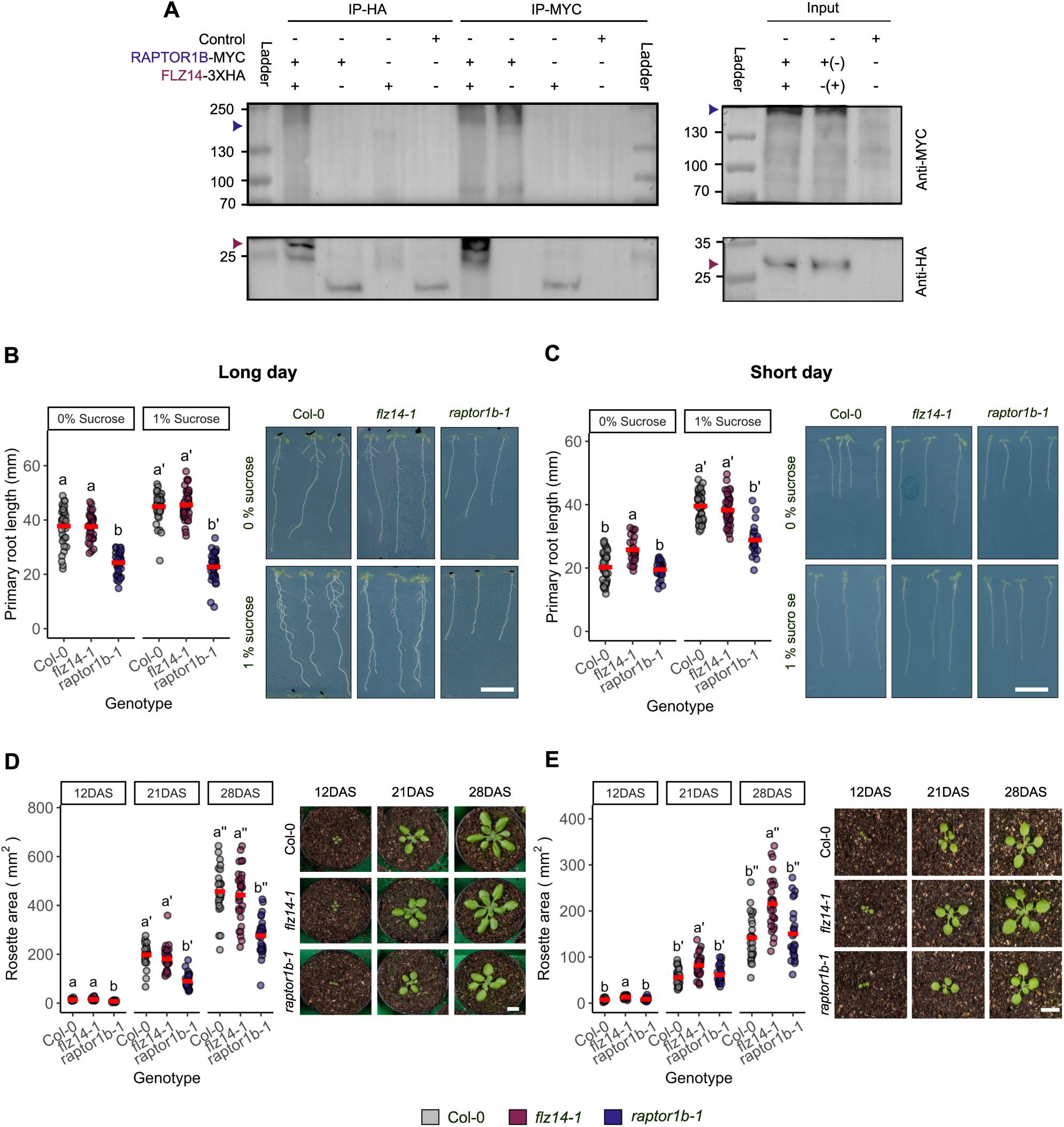
RAPTOR1B and FLZ14 physically interact and their genetic mutations trigger a photoperiod-and sugar status-dependent phenotype. **(A)** FLZ14 interacts with RAPTOR1B when transiently expressed in tobacco leaves. *35S:RAPTOR1B:6xMyc* was co-infiltrated with *35S:FLZ14-3xHA* in 5 weeks-old tobacco plants. *RAPTOR1B-6xMyc* (∼220 kDa) and *FLZ14-3xHA* (∼30 kDa) were immunoprecipitated from total protein extracts, and further immunodetected with Myc and HA-specific antibodies. The symbols + and – represent the infiltration or not of controls (empty vectors), *RAPTOR-6xMyc* and *FLZ14-3xHA*. The symbols in parentheses in the column representing the “Input” denote the vectors infiltrated and immunoblotted using HA antibody “Anti-HA”. Two independent experiments were performed leading to a similar conclusion. **(B and C)** Col-0, *flz14-1,* and *raptor1b-1* seeds were grown on ½ MS with and without sucrose (1% w/v). Ten days after stratification plates were scanned to measure primary root length using Fiji. **(B)** *raptor1b-1* presented a reduced primary root length compared to Col-0 and *flz14-1* and was insensitive to sucrose exogenous supply when grown in LD. Total number of primary roots measured in a single experiment: 47-33 per genotype: Col-0, 0% sucrose, *n =* 35, 1% sucrose, *n =* 47; *flz14-1*, 0% sucrose, *n =* 35, 1% sucrose, *n =* 45; *raptor1b-1*, 0% sucrose, *n =* 33, 1% sucrose, *n =* 44. **(C)** *flz14-1* displayed longer primary root length in 0% sucrose, but together with *raptor1b-1* has weaker growth stimulation towards 1% sucrose supply compared to Col-0 wild-type in SD. Total number of primary roots measured: 42-24 per genotype: Col-0, 0% sucrose, *n =* 39, 1% sucrose, *n =* 48; *flz14-1*, 0% sucrose, *n =* 24, 1% sucrose, *n =* 42; *raptor1b-1*, 0% sucrose, *n =* 30, 1% sucrose, *n =* 22. Two independent experiments led to similar results. **(D and E)** Col-0, *flz14-1,* and *raptor1b-1 p*lants were grown in soil for 28 days and rosette pictures were taken 12, 21, and 28 days after stratification (DAS) and further analysed on Fiji. **(D)** *flz14-1* has a wild-type-like rosette area while *raptor1b-1* exhibits a substantial growth reduction along the vegetative stage in LD. The total number of rosettes measured: 30-25 per genotype: Col-0, *n =* 25; *flz14-1*, *n =* 30; *raptor1b-1*, n = 29. Three independent experiments were performed and led to similar results. **(E)** *flz14-1* has a larger rosette along the vegetative stage in short days compared to Col-0 wild-type and *raptor1b-1* (both having similar rosette sizes) in SD. The total number of rosettes measured: 28-25 per genotype: Col-0, *n =* 27; *flz14-1*, *n =* 28; *raptor1b-1*, n = 25. Six independent experiments were performed, four of which led to similar results (one experiment did not show a mean difference between Col-0 and *flz14-1,* and one experiment did not show statistical differences between Col-0 and *flz14-1* (although the mutant exhibited larger rosette mean area). **(B-E)** A red rectangle indicates means, and letters represent the statistical difference among genotypes and per condition (within similar sucrose treatment for **(B)** and **(C)** or at similar DAS for **(D)** and **(E)**) performed using one-way analysis of variance (ANOVA) based on a linear model with a post-hoc Tukey honestly significant difference (HSD) test. The scale bars on the images of panel **(B-E)** represent 10 mm.

To further investigate the possible molecular connection between FLZ14 and RAPTOR1B, we used a mutant harbouring a T-DNA insertion in the second exon of *FLZ14*, hereafter referred to as *flz14-1* and previously characterized (Luhua et al., 2013; Yang et al., 2023), which displays a 96 % reduction in FLZ14 transcript **(Supplemental Figure 2)**. Given that *raptor1b-1* displays growth impairment under LD conditions (Anderson et al., 2005; Salem et al., 2018) and the activation of TORC activity by sugar supply (Xiong et al., 2013), we investigated the growth response of *flz14-1* and *raptor1b-1* to the photoperiod length and/or exogenous sugar supply. The root length and rosette area of *flz14-1* and wild-type Col-0 grown in LD conditions without sucrose addition were identical; in contrast, *raptor1b-1* displayed an overall reduction of 30 % in growth **(Figure 1B and Supplemental Figure 3A)**. Sucrose feeding into the media led to 18 % overall increase in growth in *flz14-1* and Col-0 seedlings but produced no effect on *raptor1b-1* **(Figure 1B and Supplemental Figure 3A)**. These results suggest that exogenous sugar supply is unable to stimulate growth in *raptor1b- 1* under longer photoperiods.

Interestingly, under SD (8h light/16h dark), the overall growth of Col-0 and *raptor1b-1* was very similar, while *flz14-1* was about 18% larger than both genotypes **(Figure 1C and Supplemental Figure 3B)**. Exogenous sucrose supply stimulated Col-0 growth by ∼48 %, reaching a similar size as *flz14-1* **(Figure 1C and Supplemental Figure 3B)**. Both *flz14-1* and *raptor1b-1* showed a smaller increase in growth than Col-0, ∼35 % more in comparison to the growth in the media lacking sucrose **(Figure 1C and Supplemental Figure 3B)**. These results suggest that *raptor1b-1* and *flz14-1* growth is dampened in response to sucrose. Furthermore, *FLZ14* mutation seems to have a larger impact on growth when sugars are less available (i.e., SD conditions).

To confirm that the *flz14-1* growth phenotype is related to the sugar-TORC network, we took advantage of the ATP-competitive TOR inhibitor AZD-8055 (Montané and Menand, 2013) to perform a sensitivity assay using this mutant. As previously reported, the root length of Col-0 plants strongly decreased as the AZD-8055 concentration increased, regardless of the photoperiod (Montané and Menand, 2013) **(Supplemental Figure 3C and D)**. While *flz14-1* root length followed the same growth pattern as Col-0 under various AZD-8055 concentrations in LD **(Supplemental Figure 3C)**, this mutant was partially insensitive to the TOR inhibitor regardless of the tested concentration in SD conditions **(Supplemental Figure 3D)**. These results further suggest that FLZ14 is part of the TORC signalling network and acts in a photoperiod-dependent manner.

To verify that the increased growth of *flz14-1* in SD was due to the loss of a functional *FLZ14*, we generated stably transformed lines expressing *FLZ14* genomic DNA driven by its native promoter and flanked with a mNeonGreen fluorescent protein at either C-terminal or N-terminal. One and five independent lines flanked with mNeonGreen at the C-terminal domain (C2) and the N-terminal domain of FLZ14 (C13, C17, C19, C20, and C21), respectively, could restore the root growth length at the same levels as the wild-type **(Supplemental Figure 3E)**.

Finally, the photoperiod-dependent growth phenotype of *flz14-1* and *raptor1b-1* was further validated in plants grown on soil. In LD, *flz14-1* rosette was identical to Col-0, while *raptor1b-1* showed smaller rosettes from 12 to 28 days after sowing (DAS) **(Figure 1D)**, confirming the results obtained *in vitro*. However, the *flz14-1* rosette was larger than Col-0 and *raptor1b-1* in SD, which were phenotypically indistinguishable from each other **(Figure 1E)**. Altogether, these point toward an antagonistic role of FLZ14 and RAPTOR1B in mediating growth, which is photoperiod-and/or sugar-dependent. While FLZ14 might repress growth under SD/low sugar conditions, RAPTOR1B stimulates growth in LD/high sugar conditions.

### FLZ14 acts as a negative regulator of TORC signalling in SD

We next speculated that TOR activity is enhanced in *flz14-1* when grown under SD. This was further assessed by measuring S6K activity, a well-known readout of the TOR activity (Xiong et al., 2013). Leaves of Col-0 and *flz14-1* grown under LD displayed similar S6K activity patterns along the diel cycle, with the only exception being at ZT8, when S6K activity slightly increased in *flz14-1* **(Figure 2A)**. Under SD, however, S6K activity was about 20% higher from dusk to dawn (ZT8 to ZT24) in *flz14*-1 compared to Col-0 **(Figure 2B)**. This agrees with the enlarged growth phenotype of *flz14-1* in this photoperiod **(Figure 1E)**, indicating that FLZ14 acts as a negative regulator of TORC signalling in SD conditions during the night.

**Figure 2.**
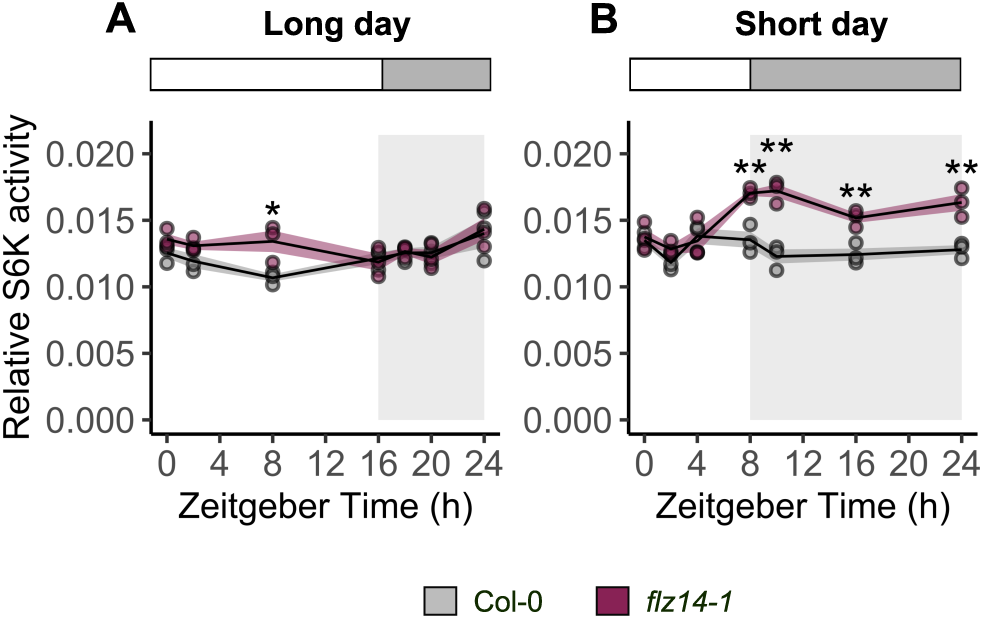
FLZ14 restricts TOR activity in SD. **(A)** *flz14-1* does not change TOR activity in LD. Col-0 and *flz14-1* were grown for 28 days in soil and whole rosettes were harvested at ZT0, 2, 8, 16, 18, 20 and 24 for assessing the phosphorylation status of S6K as a proxy for TORC activity (n = 3 biological replicates, each containing a pool of 11 rosettes). **(B)** *flz14-1* exhibits a higher TOR activity during the night in SD. Col-0 and *flz14-1* were grown for 28 days in soil and whole rosettes were harvested at ZT0, 2, 4, 8, 10, 16, and 24 for assessing the phosphorylation status of S6K as a proxy for TORC activity (n = 3 biological replicates, each containing a pool of 21 rosettes). **(A and B)** Two independent experiments led to similar results for each photoperiod. The solid black line represents the mean, and the shade-filled area represents the standard error. Asterisk indicates a significant difference between genotypes for a specific ZT (*P<0.05, **P<0.01, one-way ANOVA).

### Genetic interaction between FLZ14 and RAPTOR1B is photoperiod-dependent

To better understand the crosstalk between FLZ14 and RAPTOR1B in the regulation of growth, double mutants, namely *flz14*-*1*/*raptor1b-1,* were generated. We could confirm the similar rosette size of *flz14-1* and Col-0 in LD, which was relatively smaller in *raptor1b-1* **(Figure 3A)**. As expected, TOR activity was reduced in *raptor1b-1* in comparison to *flz14-1* and Col-0 **(Figure 3C)**. Furthermore, the *flz14-1/raptor1b-1* double mutant phenocopied *raptor1b-1* single mutant rosette size **(Figure 3A)**, both resulting in a similar decreased TOR activity with respect to Col-0 and *flz14-1* (**Figure 3C**).

**Figure 3.**
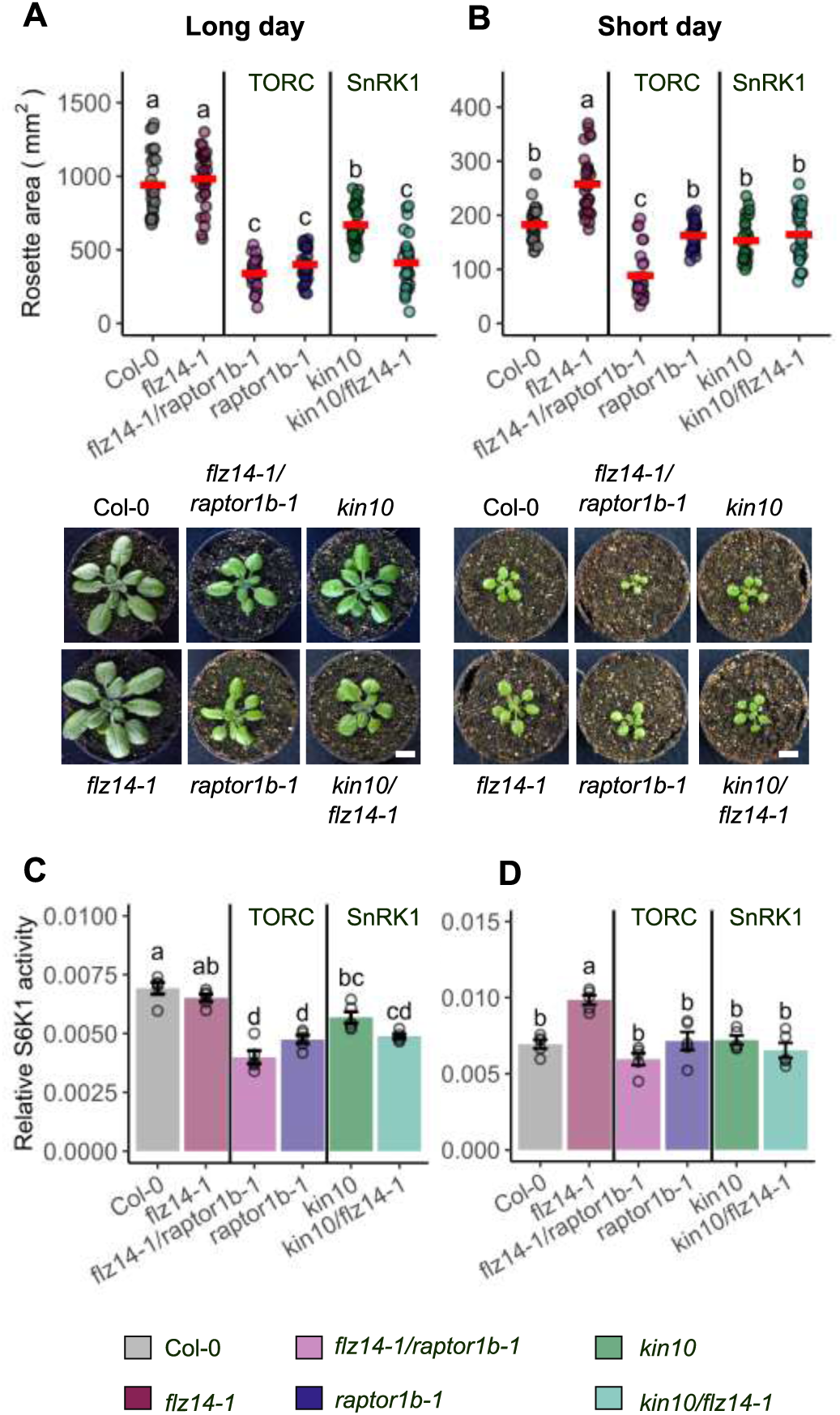
*FLZ14* genetically interacts with *RAPTOR1B* and *KIN10* in a photoperiod-dependent manner. Col-0, *flz14-1, raptor1b-1, flz14-1/raptor1b-1*, *kin10*, and *kin10/flz14-1* plants grown in LD and SD for 28 days. **(A)** Combination of *flz14* and *raptor1b* mutations results in a *raptor1b-*like rosette area (epistatic interaction), while *flz14* combined with *kin10* mutation results in a smaller rosette area compared to both parents (pleiotropic effect) in LD. Total number of rosettes measured: 30 per genotype: Col-0, *n =* 30; *flz14-1*, *n =* 30; *raptor1b-1*, n = 30; *flz14-1/raptor1b-1*, n = 30; *kin10*, *n =* 30; *kin10/flz14-1, n =* 30. **(B)** Combination of *flz14* and *raptor1b* mutations results in a smaller rosette size than both parents (pleiotropic effect) while *flz14* combined with *kin10* mutation results in a *kin10*-like rosette area (epistatic interaction) in SD. Total number of rosettes measured: 30-27 per genotype: Col-0, *n =* 30; *flz14-1*, *n =* 29; *raptor1b-1*, n = 30; *flz14-1/raptor1b-1*, n = 27; *kin10*, *n =* 27; *kin10/flz14-1, n =* 29. **(A and B)** The red rectangles indicate the means and letters represent the statistical difference between genotypes performed using one-way ANOVA based on a linear model with a post-hoc Tukey HSD test. **(C and D)** Representative images from **(A)** and **(B)**, scale 10mm. **(E)** *flz14-1/raptor1b-1*, and *kin10/flz14-1* lead to a *raptor1b-1* and *kin10-*like TOR activity, n = 5 biological replicates representing a pool of 4 rosettes. **(F)** The high TOR activity triggered by the mutation of *flz14* is suppressed in *flz14- 1/raptor1b-1*, and *kin10/flz14-1*; *n* = 5 biological replicates representing a pool of 7 rosettes. **(E and F)** Twenty-eight days old rosette of Col-0, *flz14-1, raptor1b-1, flz14-1/raptor1b-1*, *kin10*, and *kin10/flz14-1* were harvested shortly before dusk (**(E)** ZT16, **(F)** ZT 8) and total protein was extracted to assess the phosphorylation status of S6K as a proxy for TORC activity. Means are indicated by the bars fitted with error bars representing standard error, and letters denote statistical differences between genotypes based on a one-way ANOVA with a post-hoc Tukey HSD test.

Similar to the previous analysis, *flz14-1* displayed a rosette ∼ 30% bigger than *raptor1b-1* and Col-0 in SD **(Figure 3B)**. Interestingly, the *flz14-1/raptor1b-1* double mutant presented a smaller rosette area than the *raptor1b-1* and *flz14-1* single mutants **(Figure 3B)**. These results suggest that apart from being a negative regulator of the TORC mediated by RAPTOR1B, *FLZ14* might also operate partially independently from the TORC pathway at the genetic level to regulate growth in SD. This hypothesis was further verified by assessing the activity of TOR in the single and double mutants. We anticipated that TOR activity should positively correlate to growth. Indeed, this was the case for the single mutants *flz14-1 and raptor1b-1* **(Figure 3D)**. The indistinguishable growth phenotype of Col-0 and *raptor1b-1* followed by similar levels of S6K activity indicates that RAPTOR1B is less required for growth in SD.

In *flz14-1/raptor1b-1*, however, the TOR activity was similar to that in *raptor1b-1* and Col-0 but about ∼50 % lower than *flz14-1* at ZT8 **(Figure 3B)**. These results corroborate that FLZ14 operate partially independently of TORC in SD. Altogether, these data further support a conditional function of FLZ14 in regulating TOR activity through RAPTOR1B according to the sugar status and/or photoperiod.

### KIN10 is also involved in the sugar-FLZ14-TORC network

Recently, FLZ14 was shown to be involved in SnRK1-ATG genetic pathway (Yang et al., 2023). Moreover, three members of the FLZ family have been described to facilitate the interaction between SnRK1 and TORC in a condition-specific fashion (Jamsheer et al., 2018, 2022). To check whether FLZ14 operates in an SnRK1-dependent manner, we generated a *kin10/flz14-1* double mutant. In LD, *kin10* displayed a reduced rosette area **(Figure 3A)** and TOR activity **(Figure 3C)** compared to Col-0. Surprisingly, this decrease was even greater in the *kin10/flz14-1* double mutant than in both parental lines **(Figure 3A and C)**. These differences in growth cannot be solely explained by the contrasted TOR activity, implying that additional players might be involved in regulating growth under high-sugar availability conditions.

Strikingly, *kin10* and *kin10/flz14-1* rosette sizes were similar to Col-0, but smaller than *flz14- 1* in SD **(Figure 3B)**, indicating that FLZ14 depends on SnRK1 to regulate growth in SD. However, the TOR activity was unaltered in *kin10, kin10/flz14-1* with respect to Col-0, while it was enhanced in *flz14-1* **(Figure 3D)**, suggesting that the function FLZ14 and KIN10 are involved in restricting TORC activity in SD.

Our results demonstrate that FLZ14 displays a conditional function mediated by sugar fluctuations. Higher oscillation in the sugar levels, such as those triggered in SD conditions (i.e., longer nights and short days), favours the crosstalk between FLZ14 and SnRK1 to slow down the activity of TOR. In contrast, when the sugar availability is more constant, this crosstalk seems to be abolished, and TORC can fulfil its function in growth.

### FLZ14 expression is regulated by fluctuations in sugar status

To exclude the photoperiod effect on *FLZ14* regulation, we next closely inspected the relationship between its expression levels and sugar status using mutants with altered levels of endogenous sugars in plants grown under equinoctial conditions. The transcript levels of *TREHALOSE-6-PHOSPHATE SYNTHASE 5* (*TPS5)* and *DARK INDUCIBLE 6 (DIN6),* known to respond to sugar surplus and starvation, respectively, were also measured as controls (Fujiki et al., 2000; Bates et al., 2012). The analysis was focused on the transition from dark to light (ZT 22, 0, 2), when plants face large oscillations in their sugar status. To better capture the differences among the mutants, fold changes between ZT22 and ZT2 were computed. Mutation in the enzyme coding the *PLASTIC PHOSPHOGLUCOMUTASE (pgm)*, leads to acute starvation in the dark, followed by an accumulation of sugars in the light, bringing about a large fluctuation in the sugar levels (Caspar et al., 1985). Indeed, the largest *FLZ14* fold change was found in *pgm* **(Figure 4)**. This was accompanied by the highest increase in the sugar levels and the lowest fold change in the starvation marker *DIN6* **(Figure 4 and Supplemental Figure 4A)**. To a lesser extent, Col-0 displayed a similar trend than *pgm* with the only difference being the induction of *TPS5* **(Figure 4)**, confirming that *FLZ14* responds to sugar levels (Jamsheer K and Laxmi, 2015). Surprisingly, the mutant with larger endogenous sugar content, *sweet11/12* (i.e., impaired in sucrose transport), was among the mutants with the lowest *FLZ14* induction **(Figure 4)**. As expected, no differences in the sugar content fold change were noticed in this mutant, while *TPS5* expression tended to be induced at a much lesser extent than Col-0 and *pgm* **(Figure 4 and Supplemental Figure 4A)**. Taken together these data suggest that *FLZ14* expression is maximised under greater changes in sugar levels, such as the ones during the resupply of sugars to C-starved plants.

**Figure 4.**
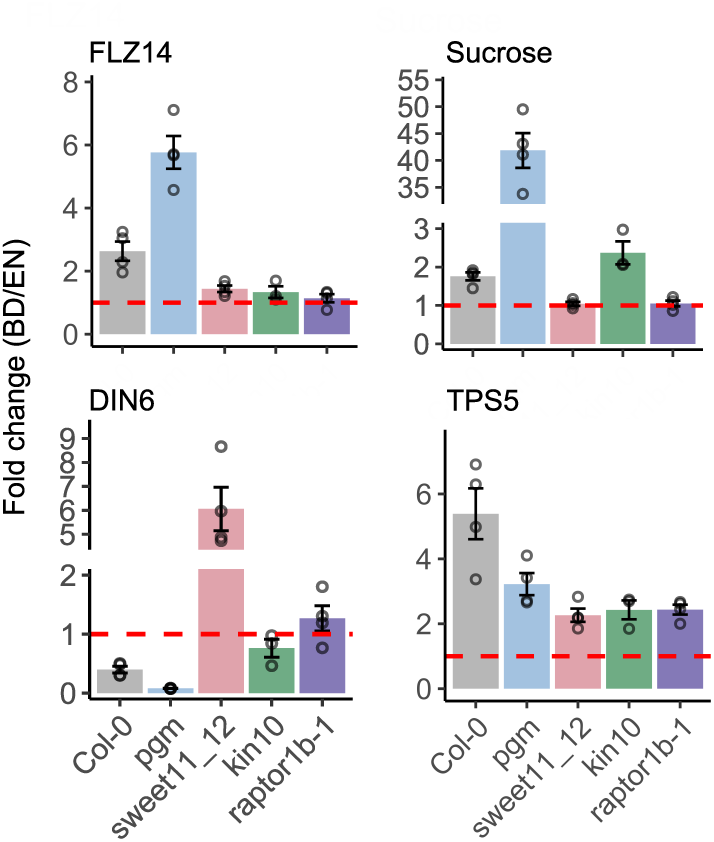
*FLZ14* expression is regulated by the difference in the sugar status between the dark and light phases. *FLZ14* expression is maximised under a larger oscillation in in sugar status. 12-days-old seedlings grown hydroponically under equinoctial days (12 h light / 12 h darkness) were harvested 2h before and after dawn referenced as the end of the night (EN) and beginning of the day (BD), respectively. Expression analysis was assessed by real-time PCR using *PP2A (AT1G13320)* and *TIP41 (AT4G34270)* as reference genes validated using NormFinder. Fold change was built between BD and EN using delta Ct or relative metabolite concentration for the expression of *FLZ14*, the starvation-marker *DIN6*, the sucrose-induced gene *TPS5*, and the content of the soluble sugar sucrose. This analysis included the following mutants with endogenous alteration in sugar status: *sweet11/12* (sugar transporter, constant and high levels of sucrose between dark and light), *pgm* (transitory starch synthesis, starchless mutant resulting in large differences in sugar content between day and night), *raptor1b-1* and *kin10* (sugar sensing pathway, altered sugar content and signalling). The fold changes (BD/EN) were performed replicate-wise for each mutant. The average of individual fold changes was plotted and fitted with error bars, representing standard error. The red dashed lines indicate a fold change equal to 1 (similar expression or sucrose content between BD and EN). n = 4 biological replicates representing a pool of 40 seedlings.

The expression of *FLZ14* and *DIN6* also did not drastically change in *raptor1b-1* and *kin10* **(Figure 4)**. In *raptor1b-1*, this trend was followed by unaltered sucrose and glucose levels, but a slight induction in *TPS5* **(Figure 4 and Supplemental Figure 4A)**. In *kin10*, all sugars were enhanced **(Figure 4 and Supplemental Figure 4A)**. Such a pattern indicates that the relationship between *FLZ14* expression and sugar status requires functional sugar signalling modules.

To further validate that the amplitude of *FLZ14* induction becomes larger as the delta sugar levels increase in the transition between night and day, we performed Spearman correlation considering all time points (ZT22, ZT0, and ZT2) **(Supplemental Figure 4B)**. The results confirm that *FLZ14* expression positively correlates with sugars and *TPS5* while negatively correlating with *DIN6*. This correlation vanishes when the fluctuation in sugar levels does not occur, as in the s*weet11/12* and *raptor1b-1* mutants **(Figure 4 and Supplemental Figure A and B)**. Altogether, this analysis showed that the regulation of *FLZ14* expression depends on the magnitude of changes in sugar status.

## Discussion

During the diel cycle, the length of the light period directly affects the production and the allocation of photoassimilates to plant growth (Gibon et al., 2009; Sulpice et al., 2009, 2014). In nature, seasonal changes and variations within a single day, cloud movement for instance, cause oscillations in photoperiod duration and sunlight intensity. Therefore, sensing fluctuations in C pools is essential for proper growth and development. TORC and SnRK1 signalling pathways are two master hubs that conveys energy/nutrients surplus and limitation signals, respectively [for reviews see (Caldana et al., 2019; Artins and Caldana, 2022; Li et al., 2021)]. A wealth of evidence supports the importance of sugars in activating the TORC pathway in plants (Xiong et al., 2013; Dobrenel et al., 2016), which in turn affects overall growth and metabolism (Moreau et al., 2012; Salem et al., 2018; Caldana et al., 2013; da Silva et al., 2021). However, most studies were based on *in vitro* exogenous sugar supply thereby overlooking the endogenous sugar fluctuations along the diel cycle. Here, we found that RAPTOR1B directly interacts with FLZ14. Changing C availability by manipulating the photoperiod length uncovered a two-way function of *FLZ14*, which favours growth regulation by TORC when C is not limiting (e.g., long days) **(Figure 5A)** while favouring SnRK1-dependent growth regulation via downregulating TORC activity in restricted C conditions (e.g., short days) **(Figure 5B)**.

**Figure 5.**
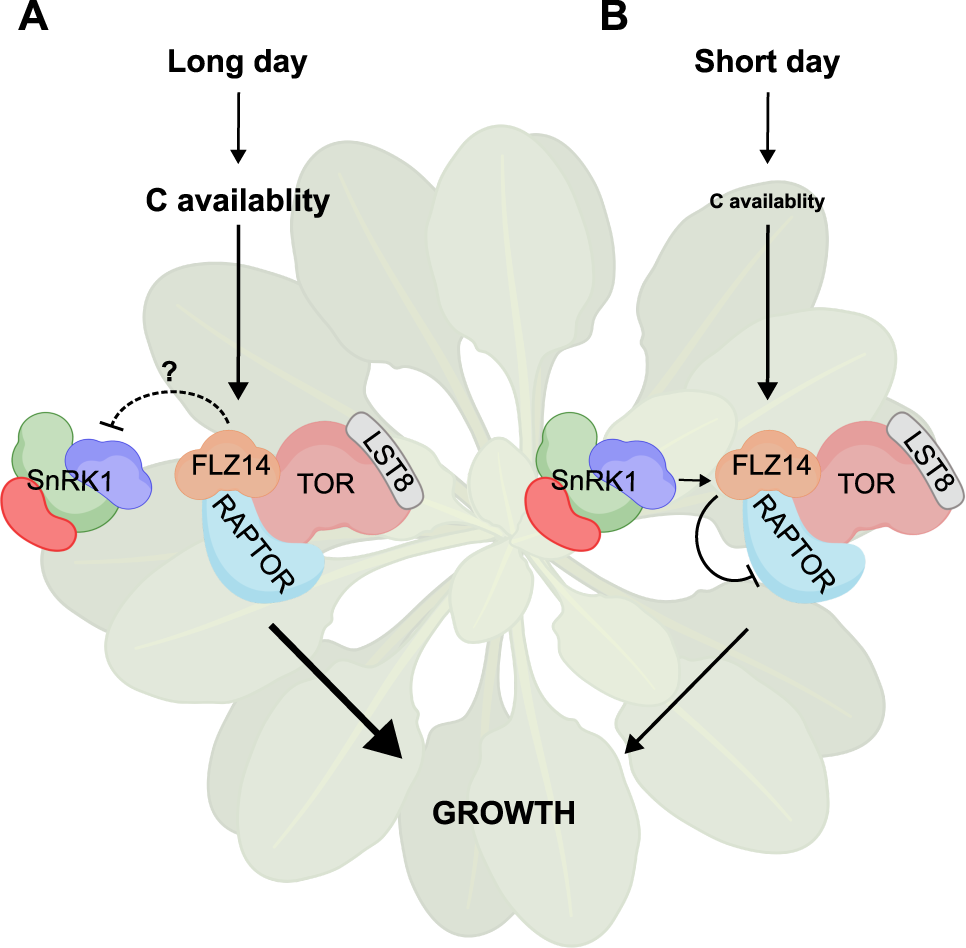
Model summarising the model of action of FLZ14 in the TORC-SnRK1 network. **(A)** Under LD conditions, FLZ14 favours TORC-dependent regulation of plant growth, which might imply the regulation of SnRK1. **(B)** Under SD conditions, FLZ14 mediates an SnRK1-dependent signal leading to the restriction of TORC activity. Arrows show a promotive effect and blunted head arrows denote a repressive effect (plain) or potential repressive effect (dashed).

Our work confirmed that *RAPTOR1B* loss-of-function mutant wrestles to grow and develop under LD conditions **(Figure 1 and 3)** (Salem et al., 2018), whereas its vegetative growth is unaffected in restricted C availability driven by SD. These findings might indicate the presence of a SD-specific TOR regulation mechanism, which is mediated by RAPTOR1B. Indeed, our data showed that FLZ14 operates as part of the network, which restricts TOR activity in SD **(Figure 2 and 3)**. First, the increased growth of *flz14-1* mutant only in SD correlates well with a higher TOR activity, especially at dusk and during the dark period **(Figure 2)**. Second, the genetic interaction between *raptor1b-1* and *flz14-1* leads to a significant reduction in growth, surpassing that of both parental lines, without compromising TOR activity **(Figure 3)**. These results indicated the need for additional players in the interplay between *RAPTOR1B* and *FLZ14* to pace growth in SD. In the LD, however, this double mutant phenocopied the growth and TOR activity of *raptor1b-1* **(Figure 3)**, suggesting that *FLZ14* might act dependently on *RAPTOR1B* to regulate the growth in LD.

Other members of FLZ family, FLZ6, FLZ10, and FLZ8, have been reported to influence TOR activity through different SnRK1-dependent mechanisms (Jamsheer et al., 2018, 2022). In the case of FLZ6 and FLZ10, this connection is indirect and dependent on KIN10 in response to stress-induced starvation (Jamsheer et al., 2018), while FLZ8 favours the interaction between RAPTOR1B and KIN10 to avoid sugar surplus-dependent TORC hyperactivation (Jamsheer et al., 2022). Our findings show that FLZ14 also interconnects KIN10 and RAPTOR1B, however, this interaction seems to be more complex. Both *kin10* single and *kin10/flz14-1* double mutants exhibit growth and TORC activity resembling the Col-0 phenotype **(Figure 3)**, implying that KIN10 alone does not repress TOR in SD. These counterintuitive results add an extra layer of complexity to this interplay as the genetic interaction between *kin10* and *flz14-1* masks the negative impact of FLZ14 on TORC/RAPTOR1B. Furthermore, the double mutant phenocopied *raptor1b* and *raptor1b/flz14-1* under LD conditions **(Figure 3)**. These findings hint at a role of FLZ14 in mediating the interplay between SnRK1 and TOR through RAPTOR1B, which is condition-dependent and probably requires other players to rewire this energy/sugar signalling network. Under unexpected C starvation (i.e., seedlings grown for 1 day of continuous light followed by 3h of darkness), SnRK1-dependent autophagy triggers degradation of FLZ14 (Yang et al., 2023). We speculate that this complexity emerged from the fine-tuned regulation required under physiological conditions used in this study. This is probably less evident in previous studies, which considered more severe stress conditions in the SnRK1-dependent repression of TOR (Nukarinen et al., 2016; Jamsheer et al., 2018, 2022; Belda-Palazón et al., 2020).

Changes in photoperiod length mitigate physiological fluctuations in sugar levels, unravelling an interconnection between TORC and SnRK1, which is more dynamic than a on/off switch relationship according to acute changes in energy/sugar status. In our work, this dynamic could be depicted by the following observations. First, genetic mutation of *KIN10* does not affect TOR activity in SD **(Figure 3)**, indicating that SnRK1 does not constantly repress TORC under C-limited conditions. Second, the reduced growth of *kin10* correlates with dampened TORC activity in LD **(Figure 3)**, raising the possibility that SnRK1 inactivates a repressor of the TOR signalling to balance growth while integrating C availability under more physiological conditions. Interestingly, a very recent study showed that FLZ14 represses SnRK1 activity under C-limiting conditions (Yang et al., 2023).

We cannot rule out that other subunits of the SnRK1 complex might play a role in this interplay, such as the second catalytic subunit of the SnRK1 complex KIN11 (Soto-Burgos and Bassham, 2017; Baena-González et al., 2007). Another possibility might be the implication of other players, such as SnRK2, which prevents SnRK1-dependent TOR repression in the absence of abscisic acid (ABA) (Belda-Palazón et al., 2020). These authors suggest that SnRK2 might regulate TORC signalling in optimal growth conditions, highlighting the complexity of this network.

Diel and seasonal changes impact greater the fluctuations in the endogenous sugar levels, influencing growth. Previous work reported the induction of *FLZ14* transcripts in response to abrupt increases in sugar levels caused either by exogenous supply (3 % glucose or sucrose) or light after prolonged darkness (Jamsheer K and Laxmi, 2015). Similarly, darkness after continuous light conditions also represses FLZ14 expression and protein levels (Yang et al., 2023). Here, we confirm that *FLZ14* expression positively correlates with the physiological levels of sugar **(Figure 4 and Supplemental Figure 4)**. However, by using mutants with change in endogenous sugar content (e.g., *pgm* and *sweet11/12*), our analysis reveals that *FLZ14* does not respond solely to high sugar status, but rather to larger oscillations in the endogenous sugar levels. *FLZ14* expression was maximised in *pgm*, which experiences sugar depletion at night but a strong accumulation of sugar shortly after the transition to light, respectively **(Figure 4 and Supplemental Figure 4)**. By contrast, *sweet11/12* keeps its sugar content high and constant throughout the diel cycle dampening *FLZ14* induction **(Figure 4 and Supplemental Figure 4)**. These findings highlight the involvement of FLZ14 in responding to significant physiological fluctuations in sugar status, such as those observed during dark/light transitions, which can also be influenced by photoperiod.

Despite the wealth of evidence supporting the need for tighter growth regulation in SD, the main players in this process remain largely unknown. Our work sheds light on this complex network by unravelling a new player: FLZ14. This protein conveys sugar status/ C availability signals to pace the TORC mediated by RAPTOR1B, a major player in growth. Furthermore, it scratches the tip of the iceberg by revealing the need for other players, such as SnRK1, to integrate metabolic signals to growth **(Figure 5)**. The elucidation of further players in the network will be of great interest to better understand the mechanism of growth regulation in plants in response to natural fluctuations in the environment.

## Methods

### Plant material and growth

Arabidopsis plants were grown in 16/8 h (long days; LD), 12/12 h (equinoctial days; ED), and 8/16 h light/dark cycles (short days; SD) at 120 μmol m^-2^ s^-1^ fluence rate, 21/19°C light/dark cycles. Single mutants were purchased from NASC, and double mutants were obtained by crossing single mutants and checked by PCR-based genotyping in the segregating T3 generation. Primers and plant lines used in this study are listed in (**Supplemental Table 2)**.

For growth in soil, single or six non-sterilised seeds were sown in 6 cm or 12 cm diameter pots, respectively, containing Arabidopsis basic medium composed mainly of white peat, stratified and transferred in SD and LD conditions. Pictures were taken 12, 21, and 28 days after stratification (DAS), and the rosette area was measured using FIJI (https://imagej.net/Fiji).

*In vitro* cultivated seeds were soaked in a bleach/EtOH solution (0.125 % / 90 %) to avoid biotic contamination, plated and stratified for 3 days at 4 °C in complete darkness. Seedlings were grown hydroponically, as previously described (Monte-Bello et al., 2018). To assess the sugars content and gene expression levels in *sweet11/12*, *pgm*, *kin10*, and *raptor1b-1*, 12 days-old seedlings grown under ED were harvested 2 and 0.5 h before, and 2 h after dawn, snapped frozen into liquid nitrogen, and stored at -80°C until processing. A total of four biological replicates representing a pool of 40 whole seedlings were used.

To assess the Arabidopsis growth *in vitro*, after stratification, two rows of 12 sterilised seeds were sown on square Petri dishes containing 80 mL of ½ MS media (Murashige and Skoog, 1962), 0.8% agar (Sigma-Aldrich, https://www.sigmaaldrich.com/DE) supplemented or not with 1% sucrose. The AZD-8055 sensitivity assay was carried out using concentrations ranging from 0 to 1 µM (LC Laboratories, https://lclabs.com/), and an equivalent volume of DMSO (solvent used to dissolve AZD-8055) (Carl Roth, https://www.carlroth.com/) was used as control. After stratification, plates were placed vertically in a growth chamber set to either SD or LD conditions. Pictures were taken 10 or 12 days after stratification with a scanner (EPSON, Perfection V800 photo), and primary root length and shoot area were measured using FIJI (https://imagej.net/Fiji).

### Plasmid construction and plant transformation

For generating the FLZ14 translational reporter lines, the mNeonGreen sequence flanked at the N and C-terminal domain by a 27 bp linker and the genomic DNA of *FLZ14* (_g_*FLZ14*) was amplified by PCR. mNeonGreen was fused at the N and C-terminal domain of FLZ14 by overlapping PCR and sequentially cloned into the *pDONR221*^TM^ *P1P2* using BP clonase^TM^. *pDONOR221* ^g*FLZ14-mNeonGreen*^ and *pDONOR221 ^mNeonGreen-^*^g*FLZ14*^ for C and N-terminal fusion, respectively, were cloned into the destination vector *pGWB501* by LR clonase^TM^. Then, the FLZ14 promoter (2.5 kb upstream of the *FLZ14* start codon) was cloned upstream of the inserted fragment by *In-Fusion^®^*using the XbaI enzyme. The primers used are listed in **(Supplemental Table 2)**. For all the cloning steps, Stellar™ *E. coli* competent cells were used (Takara, https://www.takarabio.com/). Finally, the destination vectors for the constructs above were electroporated into *A. tumefaciens* strain GV3101 for further transformation into *flz14-1* mutant with *proFLZ14:mNeonGreen-*g*FLZ14 and proFLZ14:*g*FLZ14-mNeonGreen* by floral dip.

### Yeast two-hybrid (Y2H)

Y2H screening was performed by Hybrigenics S.A.S. (Evry, France) (https://www.hybrigenics-services.com/) using the ULTImate Y2H^TM^ based cell-to-cell mating process to assess the interactions of RAPTOR1B (bait) with a cDNA library of Arabidopsis opened and closed flowers (prey). Two candidates with moderate and high confidence interactions were selected for testing 1:1 interaction with RAPTOR1B using an independent Y2H assay. For this purpose, the full-length cDNAs of *RAPTOR1B* and *FLZ14,* and a fragment from of *S6K1* (region from 9 to 879 bp downstream of the start codon was used as a positive control (Mahfouz *et al*., 2006)) were amplified by PCR, cloned into *pDONR221*^TM^ *P1P2* using BP clonase^TM^ (ThermoFisher), and sequenced using primers flanking the cDNA **(Supplemental Table 2)**. The generated constructs *pDONR221^RAPTOR1BcDNA^* and *pDONR221^FLZ14cDNA^*, *pDONR221^S6K1cDNA^* were recombined into the yeast vector harbouring either the DNA Binding Domain (DBD) at the N-terminus *(pDEST^TM^32)* or the Activation Domain (AD) at the N-terminus *(pDEST^TM^22)* (Invitrogen^TM^, USA) respectively, using LR clonase^TM^ (ThermoFisher). The constructs were further checked by sequencing using the primers listed in **(Supplemental Table 2)**. *pDEST^TM^32^RAPTOR1BcDNA^*was co-transformed either with *pDEST^TM^22*^FLZ14cDNA^ or *pDEST^TM^22^S6K1cDNA^* plasmids into the yeast strain Y2HGold (Takara Bio, USA) as previously described (Paiano et al., 2019). As negative controls, the empty vector *pDEST^TM^32 and pDEST^TM^22* were co-transformed with *pDEST^TM^22*^FLZ14cDNA^ or *pDEST^TM^22^S6K1cDNA^* and *pDEST^TM^32^RAPTOR1BcDNA^*, respectively, to check yeast auto-activation. The selection of positive transformants was carried out in double dropout media lacking Leucine and Tryptophan (-L/T), and recovered colonies were resuspended in 1X TE buffer and spotted into triple dropout media lacking Leucine, Tryptophan, and Histidine (-L/T/H) as previously described ((Paiano et al., 2019)). The constructs generated are listed in **(Supplemental Table 2)**.

### Biomolecular fluorescent complementation (BiFC)

The BiFC analysis was performed using the *pBiFC 2in1* cloning system gateway compatible (Invitrogen, http://www.invitrogen.com/) (Grefen and Blatt, 2012), allowing the cloning of the bait and the prey in the same vector. *RAPTOR1B*, *FLZ14,* and *FLZ6* (negative control for RAPTOR1B interaction(Nietzsche et al., 2016))full-length cDNAs were amplified by PCR and cloned into *pDONR221*^TM^ *P1P4* (*RAPTOR1B) or P2P3* (*FLZ14* and *FLZ6)* using BP clonase^TM^ (ThermoFisher). Sequences were confirmed by sequencing using the primers flanking cDNA listed in **(Supplemental Table 2)**. The *pDONR221^RAPTOR1BcDNA^, pDONR221^FLZ14cDNA^* and *pDONR221^FLZ6cDNA^* were subcloned into the *pBIFC* to generate *pBiFC ^nYFP-RAPTOR1BcDNA / cYFP-FLZ14cDNA^*, *pBiFC ^nYFP-RAPTOR1BcDNA / cYFP-FLZ6cDNA^*, *pBiFC ^nYFP-RAPTOR1BcDNA / FLZ14cDNA-cYFP^*, *pBiFC ^nYFP-RAPTOR1BcDNA / FLZ6cDNA-cYFP^* by LR clonase^TM^ (ThermoFisher). Sequences were confirmed by sequencing using the primers listed in **(Supplemental Table 2)**. The confirmed vectors were transformed by electroporation into *A. tumefaciens* (strain GV3101) and infiltrated in *Nicotiana benthamiana.* The imaging was performed using the TCS SP5 laser confocal scanning microscope (Leica Microsystems) 48 h after infiltration. RFP signal, used to screen transformed cells, was excited at 488 nm, and the emission was observed in a wavelength window from 578 to 598 nm. The YFP was excited at 515 nm, and the detector was set from 520 to 540 nm. The constructs generated are listed in **(Supplemental Table 2)**.

### Co-immunoprecipitation of RAPTOR1B and FLZ14 transiently expressed in tobacco leaves

*RAPTOR1B cDNA* was cloned into *pE3c* to obtain *pE3c^RAPTOR1B-6xMyc^,* using BP clonase^TM^. Both constructs were subcloned into the plant destination vector *pMDC32-HPB* using LR clonase^TM^ (obtained from Addgene, https://www.addgene.org), harbouring a 2X CaMV 35S promoter upstream of the gateway recombination sites to generate *35S:RAPTOR1B-6xMyc*. *FLZ14 cDNA* was first cloned into *pE2c* (*pE2c^FLZ14-3xHA^)* and subsequently into *pMDC32- HPB* as described for *RAPTOR1B* to obtain *pMDC32-HPB^35S:FLZ14-3xHA^*. Sequences were confirmed by sequencing using the primers listed in **(Supplemental Table 2)**. Constructs were electroporated into *A. tumefaciens* (strain GV3101) and co-infiltrated in *Nicotiana benthamiana* (Zhang et al., 2020). Tobacco leaves were harvested 48 h after the infiltration, snapped frozen into liquid nitrogen and stored at -80°C until processing. Subsequently, leaves were finely ground in liquid nitrogen, and 1 g of powder was resuspended into 4 ml of pre-cold extraction buffer [25 mM Tris-HCl pH 7.6, 15 mM magnesium chloride, 150 mM sodium chloride, 15 mM pNO_2_-PhenylPO_4_, 60 mM B-glycerophosphate, 0.1 % NO-40 (v/v), 0.1 mM sodium orthovanadate, 1 mM sodium fluoride, 1 mM PMSF, 1 µM E64, 5 % Ethyleenglycol, 100 µM MG132, 1 tablette of cOmplete^TM^,Mini, EDTA-free protease inhibitor cocktail (Sigma, https://www.sigmaaldrich.com/DE/de/product/roche/11836170001)], homogenised by vortexing and placed on ice for 20 min. Then, samples were sonicated for 15 min and centrifuged for 15 min at 14000 rpm to recover the soluble protein in the supernatant (∼4 mL). For the immunoprecipitation, 1.5 ml of extract was incubated with 50 µl of either Anti-Myc (Miltenyi Biotec, ref. 130-091-123) or Anti-HA (Miltenyi Biotec, ref. 130-091-122) microbeads for 1 h at 4 °C under constant agitation (the non-immunoprecipitated protein extract was used as input). Next, protein complexes were isolated using MACS columns (Milteny Biotec, ref. 130-042-701) following the manufacturer’s suggestion. Finally, proteins were eluted from the columns by adding pre-heated elution buffer [50 mM Tris-HCL pH 6.8, 50 mM DTT, 1% SDS, 1 mM EDTA, 0.005% bromophenol blue, 10% glycerol] at 95°C. Co-immunoprecipitated proteins were run onto 8 % acrylamide SDS PAGE gel, and proteins were blotted onto a 0.45 µm PDVF Immobilon-P membrane (Merck Millipore, ref. IPVH00010) after electrophoresis. Next, membranes were blocked using 1X TBS-T buffer (20 mM Tris base, 150 mM NaCl pH 7.6, 1mL.L^-1^ Tween20) supplemented with 5 % fat free milk for 2 h at room temperature. Overnight incubation at 4 °C with primary antibodies Anti-Myc (ThermoFisher, https://www.thermofisher.com/) and Anti-HA Sigma, https://www.sigmaaldrich.com/) diluted 1/1000 in 1X TBS-T containing 1 % fat free milk was performed. The secondary antibody Goat anti-rabbit and mouse IgG, HRP conjugate (Sigma, https://www.sigmaaldrich.com/) were used to detect the primary antibodies. Antibodies are listed in **(Supplemental Table 2)**.

### S6K activity assay

The S6K activity assay was performed following the protocol instructions with minor modifications (p70 S6K activity kit, catalogue no. ADI-EKS-470, Enzo Life Sciences). Briefly, 50 mg of leaf material was resuspended in 0.1 mL of 1X EB extraction buffer, and 30 µL was added to a p70 S6K substrate-coated ELISA microtiter plate. The reaction was initiated by adding 10 µL of ATP solution (1 µg.µL^−1^) followed by a 2 h incubation at 37°C.

The reaction was stopped by removing the supernatant from the wells, and 40 µL of phospho-specific substrate antibody was added and incubated for 1 h at room temperature. After washes, 40 µL of diluted solution of horseradish peroxidase-conjugated goat anti-rabbit IgG was added to the wells and incubated for 30 min. The washing step was repeated four times, and wells were incubated with 60 µL of tetramethylbenzidine substrate solution for another 30 min. The reaction was stopped by adding 20 µL of acid stop solution, and absorbance was measured at 450 nm using a Microplate reader from (EPOCH2 Plate Reader/Spectrophotometer, Biotek). Relative kinase activity was calculated by subtracting the sample absorbance from the blank absorbance and normalised by the volume used and the fresh weight.

### Reverse transcription-quantitative polymerase chain reaction

Fifty milligrams of plant material were used for RNA extraction using the Quick-RNA plant kit (Zymoresearch, https://www.zymoresearch.de). Then, 2 μg of RNA was used for cDNA synthesis, employing the RevertAid First Strand cDNA Synthesis kit (Thermo Fisher, https://www.thermofisher.com/) and diluted 1/20 in water. Quantification of cDNA was done by qPCR (applied biosystems 7900ht real-time PCR) with SYBR^TM^ Green master mix (Thermo Fisher, https://www.thermofisher.com/). Relative expression was calculated based on the delta CT method (Livak and Schmittgen, 2001)using *PP2A* (*At1g13320*) and *TIP41* (*At4g34270*) as reference genes (both were previously validated using the R script ""NormFinder"" per genotype and conditions) (De Spiegelaere et al., 2015). The primers used are listed in **(Supplemental Table 2).**

### Primary metabolite extraction and quantification

Metabolites were extracted from 50 mg of frozen ground plant material resuspended in 1mL pre-cooled MTBE extraction buffer followed by a phase separation by adding 0.5 mL of water: methanol (3:1) (v/v) (Giavalisco et al., 2011). The aqueous phase containing primary metabolites was concentrated, derivatised using *N-methyl-N-trimethylsilyltrifluoroacetamid*, and processed by gas chromatography (GC) (7890N and Agilent) using the time of flight coupled with a mass spectrometry (Pegasus HT and Leco) (GC-TOF-MS) (Lisec et al., 2006). The detection of peaks, retention time alignment and metabolite matching to the library were performed using the Target Search R package (Cuadros-Inostroza et al., 2009). Metabolite quantification based on the peak intensity was normalised by the sample fresh weight (FW) and the total ion count followed by a scaling based on log_2_ transformation for the representation.

### Statistical analysis

R studio version 2021.09.0 was used to visualise, manipulate, plot, and analyse the data. Plots were generated using the R package “ggplot<u>2</u>”, and the analysis of variance (ANOVA) using the package “<u>agricolae</u>”. All statistical parameters, including *n* number, computation of means, statistical test are described in the figure legends.

## Acknowledgements

We thank Z. Nikoloski for advising on the correlation analysis, A. Leisse for the metabolite measurements, the Greenteam for excellent plant care, the Nottingham Arabidopsis stock center for T-DNA mutant seeds. This work was supported by the Max Planck Society. We apologise in advance to colleagues whose work we were not able to cover in this manuscript.

## Author contributions

A.A. designed and performed the experiments, analyzed and interpreted the data. R.U.C conducted the yeast 2-hybrid and the co-immunoprecipitation assays. M.M.S contributed to gene expression analysis of the mutants with alteration in endogenous sugar levels which was suggested by A.R.F. A.R.F and T.A.M. critically read the manuscript. A.S for introducing the concept of sugar homeostasis in a fluctuating environment. C.C. conceived the project, supervised the research and interpreted the data. A.A. and C.C. prepared the figures and wrote the manuscript. All authors read and approved the manuscript.

---

**Supplemental Figure 1. FLZ14 physically interacts with RAPTOR1B.**

**(A)** Yeast two-hybrid assay shows the physical interaction between RAPTOR1B and FLZ14, or RAPTOR1B and S6K1 (positive control). SD2 and SD3 indicate the media lacking Leucine and Tryptophan or Leucine, Tryptophan and Histidine, respectively. FLZ14/-, S6K1/-, and -/RAPTOR1B are the yeast cells transformed solely with the prey or the bait to test the auto-activation. Slight auto-activation was found in non-diluted -/RAPTOR1B (annotaded 1), but faded in the diluted yeast (annotated 10^-1^). Three independent experiments showed similar results, however, a single experiment in represented.

**(B)** Bimolecular fluorescent complementation assay showing the interaction between RAPTOR1B and FLZ14 in tobacco leaves using both combinations *nYFP-RAPTOR1B_cDNA_/ FLZ14_cDNA_-cYFP* (middle panel) or *nYFP-RAPTOR1B_cDNA_/ cYFP-FLZ14_cDNA_* (bottom panel) all driven by the Cauliflower Mosaic virus 35S promoter. *n* and *c* refer to the N-or C-terminal region, respectively, in which the marker gene was flanked to the target protein (FLZ14 or RAPTOR1B). *nYFP-RAPTOR1B_cDNA_/FLZ6_cDNA_-cYFP* were used as a negative control. The red channel indicates the transformed cells. Three independent experiments were performed, and similar results were observed in two of them.

**Supplemental Figure 2. Validation of the T-DNA insertion line into *FLZ14* gene. Upper panel.**

Scheme showing the insertion site of the *flz14-1* T-DNA mutant (triangle) in the second exon and the qPCR primers used to check *FLZ14* expression in this study. **Lower panel,** Expression of *FLZ14* in Col-0 (grey) and *flz14-1* (purple) revealing a *knock-down* (4 % of residual expression) by qPCR. Bars represent the average, and error bars represent se. *PP2A* and *TIP41* were used as housekeeping genes.

**Supplemental Figure 3. Characterisation and functional complementation of *flz14-1* mutant.**

**(A)** In the presence or absence of sucrose, *flz14-1* mutants exhibit wild-type plant-like rosette area phenotypes, whereas *raptor1b-1* mutants display reduced rosette sizes when grown in LD. Total number of rosettes measured: 40-16 per genotype: Col-0, 0% sucrose, *n =* 24, 1% sucrose, *n =* 27; *flz14-1*, 0% sucrose, *n =* 24, 1% sucrose, *n =* 29; *raptor1b-1*, 0% sucrose, *n =* 22, 1% sucrose, *n =* 23.

**(B)** *flz14-1* and *raptor1b-1* have a decreased sensitivity to sucrose when grown under SD conditions. Total number of rosettes measured: 40-16 per genotype: Col-0, 0% sucrose, *n =* 30, 1% sucrose, *n =* 38; *flz14-1*, 0% sucrose, *n =* 24, 1% sucrose, *n =* 40; *raptor1b-1*, 0% sucrose, *n =* 16, 1% sucrose, *n =* 21.

**(A and B)** Single experiment was performed. Col-0, *flz14-1,* and *raptor1b-1* were grown on ½ MS with and without sucrose (1% w/v). Ten days after stratification, rosettes were separated from the root and arranged on a flat surface in preparation for scanning. Rosette area was measured using Fiji. The red rectangles indicate the means, and the letters represent the statistical difference between genotypes performed using one-way ANOVA based on a linear model with a post-hoc Tukey HSD test.

**(C)** *flz14-1* has a wild-type like AZD-8055 sensitivity under LD. Col-0 and *flz14-1* seeds were sown on ½ MS media containing 1% sucrose supplemented with AZD-8055 ranging from 0 (DMSO mock) to 1 µM. Total number of primary roots measured: 48-30 per genotype: Col-0, 0 µM, *n =* 34, 0.2 µM, *n =* 39, 0.4 µM, *n =* 40, 0.6 µM, *n =* 36, 0.8 µM, *n =* 37, 1 µM, *n =* 39, *flz14-1*, 0 µM, *n =* 36, 0.2 µM, *n =* 33, 0.4 µM, *n =* 39, 0.6 µM, *n =* 36, 0.8 µM, *n =* 40, 1 µM, *n =* 37.

**(D)** *flz14-1* exhibits a decreased sensitivity to AZD-8055 compared wild-type plants in SD. Col-0 and *flz14-1* seeds were sown on ½ MS media containing 1% sucrose supplemented with AZD-8055 ranging from 0 (DMSO mock) to 1 µM. Due to the sucrose-rescued *flz14-1* phenotype, exogenous sucrose was applied for the AZD-8055 sensitivity assay to avoid masking-effect of the root length phenotype in the mutant. Total number of primary roots measured: 48-30 per genotype: Col-0, 0 µM, *n =* 48, 0.2 µM, *n =* 36, 0.4 µM, *n =* 47, 0.6 µM, *n =* 36, 0.8 µM, *n =* 39, 1 µM, *n =* 37, *flz14-1*, 0 µM, *n =* 42, 0.2 µM, *n =* 30, 0.4 µM, *n =* 40, 0.6 µM, *n =* 38, 0.8 µM, *n =* 37, 1 µM, *n =* 38.

(C and D) A single experiment was performed. The red rectangles indicate the means, and the stars denote statistical difference between for genotypes performed a given treatment ZT (***P<0.001, ****P<0.0001, one-way ANOVA).

**(E)** Primary root length of the FLZ14 complemented lines 10 DAS. C1-C12 and C13-C21 refers to C-terminal and N-terminal mNeongreen fusion, respectively. The red rectangles indicate the means and the letters represent the statistical difference between genotypes performed using one-way ANOVA based on a linear model with a post-hoc Tukey HSD test.

**Supplemental Figure 4. Regulation of *FLZ14* expression is influenced by sugar status.**

**(A)** Fold change was built between BD and EN using the relative metabolite content of fructose and glucose. 12-days-old seedlings grown hydroponically under equinoctial days (12 h light / 12 h darkness) were harvested at 2 h before and after dawn referenced as the end of the night (EN) and beginning of the day (BD), respectively. This analysis included the following mutants having internal altered sugar status: *sweet11/12* (sugar transporter, constant and high levels of sucrose between dark and light, *pgm* (transitory starch synthesis, starchless mutant resulting in large differences in sugar content between day and night), *raptor1b-1* and *kin10* (sugar sensing pathway, altered sugar content and signalling). The fold changes (BD/EN) were performed replicate-wise for each mutant and individual fold changes were averaged, represented by the bars fitted with error bars representing standard error. The red dashed lines indicate a fold change equal to 1 (similar expression or sucrose content between BD and EN).

**(B)** Correlation analysis was carried out between the expression of *FLZ14, DIN6, TPS5* and the content of sucrose, glucose and fructose. The values (*R*) represent the Spearman correlation coefficient, and the stars (*P*) represent the *P-value*. *R* and *P* were performed genotype-wise. 12-days-old seedlings grown hydroponically under equinoctial days (12 h light / 12 h darkness) were harvested at 2 and 0.5 h before dawn or 2 h after dawn.

**(A and B)** Four biological replicates were used, and 40 seedlings were pooled per biological replicates.

## Bibliography

Anderson, G.H., Veit, B., and Hanson, M.R. (2005). The Arabidopsis AtRaptor genes are essential for post-embryonic plant growth. BMC Biol. 3: 1–11.

Apelt, F., Breuer, D., Olas, J.J., Annunziata, M.G., Flis, A., Nikoloski, Z., Kragler, F., and Stitt, M. (2017). Circadian, Carbon, and Light Control of Expansion Growth and Leaf Movement. Plant Physiol. 174: 1949–1968.

Artins, A. and Caldana, C. (2022). The metabolic homeostaTOR: The balance of holding on or letting grow. Curr. Opin. Plant Biol. 66: 102196.

Baena-González, E., Rolland, F., Thevelein, J.M., and Sheen, J. (2007). A central integrator of transcription networks in plant stress and energy signalling. Nature 448: 938–942.

Bates, G.W., Rosenthal, D.M., Sun, J., Chattopadhyay, M., Peffer, E., Yang, J., Ort, D.R., and Jones, A.M. (2012). A Comparative Study of the Arabidopsis thaliana Guard-Cell Transcriptome and Its Modulation by Sucrose. PLoS One 7: e49641.

Belda-Palazón, B., Adamo, M., Valerio, C., Ferreira, L.J., Confraria, A., Reis-Barata, D., Rodrigues, A., Meyer, C., Rodriguez, P.L., and Baena-González, E. (2020). A dual function of SnRK2 kinases in the regulation of SnRK1 and plant growth. Nat. Plants 2020 611 6: 1345–1353.

Caldana, C., Li, Y., Leisse, A., Zhang, Y., Bartholomaeus, L., Fernie, A.R., Willmitzer, L., and Giavalisco, P. (2013). Systemic analysis of inducible target of rapamycin mutants reveal a general metabolic switch controlling growth in Arabidopsis thaliana. Plant J. 73: 897–909.

Caldana, C., Martins, M.C.M., Mubeen, U., and Urrea-Castellanos, R. (2019). The magic ‘hammer’ of TOR: the multiple faces of a single pathway in the metabolic regulation of plant growth and development. J. Exp. Bot. 70: 2217–2225.

Caspar, T., Huber, S.C., and Somerville, C. (1985). Alterations in Growth, Photosynthesis, and Respiration in a Starchless Mutant of Arabidopsis thaliana (L.) Deficient in Chloroplast Phosphoglucomutase Activity. Plant Physiol. 79: 11–17.

Cuadros-Inostroza, Á., Caldana, C., Redestig, H., Kusano, M., Lisec, J., Peña-Cortés, H., Willmitzer, L., and Hannah, M.A. (2009). TargetSearch - a Bioconductor package for the efficient preprocessing of GC-MS metabolite profiling data. BMC Bioinformatics 10: 1–12.

Dobrenel, T. et al. (2016). The Arabidopsis TOR Kinase Specifically Regulates the Expression of Nuclear Genes Coding for Plastidic Ribosomal Proteins and the Phosphorylation of the Cytosolic Ribosomal Protein S6. Front. Plant Sci. 7: 1611.

Fujiki, Y., Ito, M., Nishida, I., and Watanabe, A. (2000). Multiple Signaling Pathways in Gene Expression during Sugar Starvation. Pharmacological Analysis of din Gene Expression in Suspension-Cultured Cells of Arabidopsis. Plant Physiol. 124: 1139– 1148.

Giavalisco, P., Li, Y., Matthes, A., Eckhardt, A., Hubberten, H.-M., Hesse, H., Segu, S., Hummel, J., Köhl, K., and Willmitzer, L. (2011). Elemental formula annotation of polar and lipophilic metabolites using 13C, 15N and 34S isotope labelling, in combination with high-resolution mass spectrometry. Plant J. 68: 364–376.

Gibon, Y., Bläsing, O.E., Palacios-Rojas, N., Pankovic, D., Hendriks, J.H.M., Fisahn, J., Höhne, M., Günther, M., and Stitt, M. (2004). Adjustment of diurnal starch turnover to short days: depletion of sugar during the night leads to a temporary inhibition of carbohydrate utilization, accumulation of sugars and post-translational activation of ADP-glucose pyrophosphorylase in the followin. Plant J. 39: 847–862.

Gibon, Y., Pyl, E.T., Sulpice, R., Lunn, J.E., HÖhne, M., GÜnther, M., and Stitt, M. (2009). Adjustment of growth, starch turnover, protein content and central metabolism to a decrease of the carbon supply when Arabidopsis is grown in very short photoperiods. Plant, Cell Environ. 32: 859–874.

Grefen, C. and Blatt, M.R. (2012). A 2in1 cloning system enables ratiometric bimolecular fluorescence complementation (rBiFC). Biotechniques 53: 311–314.

Hara, K., Maruki, Y., Long, X., Yoshino, K. ichi, Oshiro, N., Hidayat, S., Tokunaga, C., Avruch, J., and Yonezawa, K. (2002). Raptor, a Binding Partner of Target of Rapamycin (TOR), Mediates TOR Action. Cell 110: 177–189.

Jamsheer K, M. and Laxmi, A. (2015). Expression of Arabidopsis FCS-Like Zinc finger genes is differentially regulated by sugars, cellular energy level, and abiotic stress. Front. Plant Sci. 6: 746.

Jamsheer, M.K., Jindal, S., Sharma, M., Awasthi, P. S S., Sharma, M., Mannully, C.T., and Laxmi, A. (2022). A negative feedback loop of TOR signaling balances growth and stress-response trade-offs in plants. Cell Rep. 39: 110631.

Jamsheer, M.K. and Laxmi, A. (2014). DUF581 is plant specific FCS-like zinc finger involved in protein-protein interaction. PLoS One 9.

Jamsheer, M.K., Manvi, S., Dhriti, S., Chanchal Thomas, M., Sunita, J., Brihaspati N, S., and Laxmi, A. (2018). FCS-like zinc finger 6 and 10 repress SnRK1 signalling in Arabidopsis. Plant J. 94: 232–245.

Kim, D.H., Sarbassov, D.D., Ali, S.M., King, J.E., Latek, R.R., Erdjument-Bromage, H., Tempst, P., and Sabatini, D.M. (2002). mTOR Interacts with Raptor to Form a Nutrient-Sensitive Complex that Signals to the Cell Growth Machinery. Cell 110: 163– 175.

Li, L., Liu, K., and Sheen, J. (2021). Dynamic Nutrient Signaling Networks in Plants. https://doi.org/10.1146/annurev-cellbio-010521-015047 37: 20–21.

Lisec, J., Schauer, N., Kopka, J., Willmitzer, L., and Fernie, A.R. (2006). Gas chromatography mass spectrometry–based metabolite profiling in plants. Nat. Protoc. 2006 11 1: 387–396.

Livak, K.J. and Schmittgen, T.D. (2001). Analysis of Relative Gene Expression Data Using Real-Time Quantitative PCR and the 2−ΔΔCT Method. Methods 25: 402–408.

Luhua, S. et al. (2013). Linking genes of unknown function with abiotic stress responses by high-throughput phenotype screening. Physiol. Plant. 148: 322–333.

Mahfouz, M.M., Kim, S., Delauney, A.J., and Verma, D.P.S. (2006). Arabidopsis TARGET OF RAPAMYCIN Interacts with RAPTOR, Which Regulates the Activity of S6 Kinase in Response to Osmotic Stress Signals. Plant Cell 18: 477–490.

Mengin, V., Pyl, E.T., Moraes, T.A., Sulpice, R., Krohn, N., Encke, B., and Stitt, M. (2017). Photosynthate partitioning to starch in arabidopsis thaliana is insensitive to light intensity but sensitive to photoperiod due to a restriction on growth in the light in short photoperiods. Plant Cell Environ. 40: 2608–2627.

Montané, M.-H. and Menand, B. (2013). ATP-competitive mTOR kinase inhibitors delay plant growth by triggering early differentiation of meristematic cells but no developmental patterning change. J. Exp. Bot. 64: 4361–4374.

Monte-Bello, C.C., Araujo, E.F., Martins, M.C.M., Mafra, V., Silva, V.C.H. da Celente, V., and Caldana, C. (2018). A Flexible Low Cost Hydroponic System for Assessing Plant Responses to Small Molecules in Sterile Conditions. JoVE (Journal Vis. Exp. 2018: e57800.

Moreau, M., Azzopardi, M., Clément, G., Dobrenel, T., Marchive, C., Renne, C., Martin-Magniette, M.-L., Taconnat, L., Renou, J.-P., Robaglia, C., and Meyer, C. (2012). Mutations in the Arabidopsis homolog of LST8/GβL, a partner of the target of Rapamycin kinase, impair plant growth, flowering, and metabolic adaptation to long days. Plant Cell 24: 463–81.

Nietzsche, M., Landgraf, R., Tohge, T., and Börnke, F. (2016). A protein–protein interaction network linking the energy-sensor kinase SnRK1 to multiple signaling pathways in Arabidopsis thaliana. Curr. Plant Biol. 5: 36–44.

Nietzsche, M., Schießl, I., and Börnke, F. (2014). The complex becomes more complex: protein-protein interactions of SnRK1 with DUF581 family proteins provide a framework for cell-and stimulus type-specific SnRK1 signaling in plants. Front. Plant Sci. 5: 1–13.

Nojima, H., Tokunaga, C., Eguchi, S., Oshiro, N., Hidayat, S., Yoshino, K.I., Hara, K., Tanaka, N., Avruch, J., and Yonezawa, K. (2003). The Mammalian Target of Rapamycin (mTOR) Partner, Raptor, Binds the mTOR Substrates p70 S6 Kinase and 4E-BP1 through Their TOR Signaling (TOS) Motif. J. Biol. Chem. 278: 15461–15464.

Nukarinen, E., Nägele, T., Pedrotti, L., Wurzinger, B., Mair, A., Landgraf, R., Börnke, F., Hanson, J., Teige, M., Baena-Gonzalez, E., Dröge-Laser, W., and Weckwerth, W. (2016). Quantitative phosphoproteomics reveals the role of the AMPK plant ortholog SnRK1 as a metabolic master regulator under energy deprivation. Sci. Rep. 6: 1–19.

Paiano, A., Margiotta, A., Luca, M. De, and Bucci, C. (2019). Yeast Two-Hybrid Assay to Identify Interacting Proteins. Curr. Protoc. Protein Sci. 95: e70.

Salem, M.A., Li, Y., Bajdzienko, K., Fisahn, J., Watanabe, M., Hoefgen, R., Schöttler, M.A., and Giavalisco, P. (2018). RAPTOR Controls Developmental Growth Transitions by Altering the Hormonal and Metabolic Balance. Plant Physiol. 177: 565– 593.

Schalm, S.S., Fingar, D.C., Sabatini, D.M., and Blenis, J. (2003). TOS Motif-Mediated Raptor Binding Regulates 4E-BP1 Multisite Phosphorylation and Function. Curr. Biol. 13: 797–806.

da Silva, V.C.H., Martins, M.C.M., Calderan-Rodrigues, M.J., Artins, A., Monte Bello, C.C., Gupta, S., Sobreira, T.J.P., Riaño-Pachón, D.M., Mafra, V., and Caldana, C. (2021). Shedding Light on the Dynamic Role of the “Target of Rapamycin” Kinase in the Fast-Growing C4 Species Setaria viridis, a Suitable Model for Biomass Crops. Front. Plant Sci. 12.

Soto-Burgos, J. and Bassham, D.C. (2017). SnRK1 activates autophagy via the TOR signaling pathway in Arabidopsis thaliana. PLoS One 12: e0182591.

De Spiegelaere, W., Dern-Wieloch, J., Weigel, R., Schumacher, V., Schorle, H., Nettersheim, D., Bergmann, M., Brehm, R., Kliesch, S., Vandekerckhove, L., and Fink, C. (2015). Reference gene validation for RT-qPCR, a note on different available software packages. PLoS One 10: e0122515.

Sulpice, R. et al. (2009). Starch as a major integrator in the regulation of plant growth. Proc. Natl. Acad. Sci. 106: 10348–10353.

Sulpice, R., Flis, A., Ivakov, A.A., Apelt, F., Krohn, N., Encke, B., Abel, C., Feil, R., Lunn, J.E., and Stitt, M. (2014). Arabidopsis Coordinates the Diurnal Regulation of Carbon Allocation and Growth across a Wide Range of Photoperiods. Mol. Plant 7: 137–155.

Xiong, Y., McCormack, M., Li, L., Hall, Q., Xiang, C., and Sheen, J. (2013). Glucose-TOR signalling reprograms the transcriptome and activates meristems. Nature 496: 181– 186.

Yang, C., Li, X., Yang, L., Chen, S., Liao, J., Li, K., Zhou, J., Shen, W., Zhuang, X., Bai, M., Bassham, D.C., and Gao, C. (2023). A Positive Feedback Regulation of SnRK1 Signaling by Autophagy in Plants. Mol. Plant 16: 1192–1211.

Yazdanbakhsh, N., Sulpice, R., Graf, A., Stitt, M., and Fisahn, J. (2011). Circadian control of root elongation and C partitioning in Arabidopsis thaliana. Plant, Cell Environ. 34: 877–894.

Zhang, Y., Chen, M., Siemiatkowska, B., Toleco, M.R., Jing, Y., Strotmann, V., Zhang, J., Stahl, Y., and Fernie, A.R. (2020). A Highly Efficient Agrobacterium -Mediated Method for Transient Gene Expression and Functional Studies in Multipe Plant Species. Plant Commun.: 100028.

